# DirectMS1Quant: ultrafast quantitative proteomics with MS/MS-free mass spectrometry

**DOI:** 10.1101/2022.05.13.489895

**Authors:** Mark V. Ivanov, Julia A. Bubis, Vladimir Gorshkov, Irina A. Tarasova, Lev I. Levitsky, Elizaveta M. Solovyeva, Anastasiya V. Lipatova, Frank Kjeldsen, Mikhail V. Gorshkov

**Affiliations:** V. L. Talrose Institute for Energy Problems of Chemical Physics, N. N. Semenov Federal Research Center of Chemical Physics, Russian Academy of Sciences, 119334 Moscow, Russia; Department of Biochemistry and Molecular Biology, University of Southern Denmark, DK-5230, Odense M, Denmark; Engelhardt Institute of Molecular Biology, Russian Academy of Sciences, 119991 Moscow, Russia

## Abstract

Recently, we presented the DirectMS1 method of ultrafast proteome-wide analysis based on minute-long LC gradients and MS1-only mass spectra acquisition. Currently, the method provides the depth of human cell proteome coverage of 2500 proteins at 1% false discovery rate (FDR) when using 5-min LC gradients and 7.3 min runtime in total. While the standard MS/MS approaches provide 4000 to 5000 protein identifications within a couple of hours of instrumentation time, we advocate here that the higher number of identified proteins does not always translate into better quantitation quality of the proteome analysis. To further elaborate on this issue we performed one-by-one comparison of quantitation results obtained using DirectMS1 with three popular MS/MS-based quantitation methods: label-free quantification (LFQ), tandem mass tag (TMT), both based on data dependent acquisition (DDA), and data independent acquisition (DIA). For the comparison we performed a series of proteome-wide analysis of well-characterized (ground truth) and biological relevant samples, including a mix of UPS1 proteins spiked at different concentrations into *E. coli* digest used as a background and a set of glioblastoma cell lines. MS1-only data was analyzed using a novel quantitation workflow called DirectMS1Quant developed in this work. The results obtained in this study demonstrated comparable quantitation efficiency of 5 min DirectMS1 with both TMT and DIA methods utilizing 10 to 20-fold longer instrumentation time.

## Introduction

Personalized medicine, clinical studies, drug discoveries and many other areas of biomedical research rely on proteome analysis based on liquid chromatography in combination with high resolution tandem mass spectrometry (LC-MS/MS). Typically, this analysis employs the so-called bottom-up approach^1^ based on two possible methods: data dependent acquisition^2^ (DDA) and data independent acquisition^3^ (DIA) or the combination of both^4^. Recent advances in high resolution mass-spectrometry resulted in dramatically increased depth, throughput, and sensitivity of proteome coverage with 4000 to 5000 proteins being identified routinely within a couple of hours of total analysis time and a striking recent demonstration of up to 10,000 proteins identified in 100 min for human cell proteomes using high-resolution Orbitrap mass spectrometry^5^. In spite of these achievements, a growing trend in recent developments of technologies used for proteome analysis is reducing its time^6–13^. Indeed, the widely used DDA method involves sequential isolation of eluting peptides followed by their fragmentation to produce sequence specific MS/MS spectra. Due to the proteome digest complexity, the throughput of the MS/MS-based analysis is challenged by the speed of the high resolution mass analyzer and the peak capacity of the separation system. The fragmentation spectra are only produced for a small fraction of all ionized peptides from the sample, and this fraction is rapidly squeezed further, when the experimental acquisition is reduced to a minute time scale. The result of such a squeeze is that fewer peptides are identified for each protein with the negative effect on the quantitation accuracy, as well as a bias toward identification of larger and/or more abundant proteins. Among the possible approaches to increase the throughput of proteome-wide analysis are the so-called MS/MS-free data acquisition methods^8,14,15^ having its roots from peptide mass fingerprint approaches^16–19^. One of them, called DirectMS1 method, employs MS1-only data acquisition using high resolution mass analyzers and identifies proteins directly in the (*m/z*; retention time) search space^6^. At the core of the method is the search engine ms1searchpy^8^ based on state-of-the-art machine learning algorithms and tools for peptide feature detection in MS1 spectra and retention time prediction. Because the method is truly MS/MS-free it allows the use of ultrashort LC gradients squeezing the whole proteome analysis to minutes, yet preserving the meaningful depth. For example, when extending the peptide separation space further by employing ion mobility data, the method demonstrated identification of more than 2000 proteins of human cell proteome with 5 min LC gradients and total analysis time of 7.3 min. While this proteome coverage depth is comparable with the ones reported recently^13^ using rapid DIA-based approaches, the important feature of DirectMS1 method is high average sequence coverage of identified proteins that results in improved quantitation compared with MS/MS-based analyses.

Note that most studies on the characterization of the whole proteomes are usually focused on reporting the number of quantified proteins, omitting the more important question of quantitation efficiency or accuracy. Indeed, one of the common perceptions is that a higher number of proteins with lower CV quantitation values based on MS1 or MS2 intensities means a better quantitation method^12,13,20^. However, we advocate here that the high number of identified proteins does not always translate into better quantitation results. For example, due to higher sequence coverage for identified proteins obtained by DirectMS1, the quantitation results are better compared with MS/MS-based analysis when using short LC gradients despite the false discovery rate (FDR) being significantly compromised at the peptide level for the former method. Only a few of the protein quantitation methods account for the peptide identification quality in the quantitative results at the protein level, such as Triqler^21,22^.

In the further efforts to develop an ultra-fast MS/MS-free method of proteome-wide analysis we introduce here a novel quantitation method called DirectMS1Quant, which allows transforming peptide identifications in MS1 spectra with higher false positive rates into protein quantitation results of sound quality. To demonstrate the quantitation efficiency of DirectMS1Quant we performed proteome-wide analysis using DirectMS1 and a number of widely employed MS/MS-based methods for the well-characterized (ground truth) and real world proteomic samples.

## Experimental section

### Samples

#### Glioblastoma multiforme (GBM) samples

Label-free LC-MS/MS experiments with two low-passage cultures of GBM (3821, 4114) and normal astrocytes (AN) cell lines were used from previous study^23^ (ProteomeXchange id PXD022906). Briefly, cell cultures were grown in four replicates and treated with recombinant interferon (IFN) α-2b (4 control, 4 treated samples). Cells were resuspended in 100 µL of lysis buffer (0.1% w/v ProteaseMAX Surfactant (Promega, Madison, WI, USA) in 50 mM ammonium bicarbonate, and 10% v/v acetonitrile (ACN)) and stirred for 60 min at 1000 rpm at room temperature. Lysis was performed by sonication for 5 min at 30% amplitude on ice (Bandelin Sonopuls HD2070, Berlin, Germany). Protein extracts were reduced in 10 mM dithiothreitol (DTT) at 56 °C for 20 min and alkylated in 10 mM iodoacetamide (IAA) at room temperature for 30 min in the dark. Then, samples were overnight digested at 37 °C using sequencing grade modified trypsin (Promega, Madison, WI, USA) added at the ratio of 1:50 w/w. Digestion was terminated by acetic acid (5% w/v). Peptides were desalted using cartridges for solid-phase extraction (Oasis HLB, 1 cc, 30 mg, 30 µm particle size, Waters, Milfold, CT, USA) and dried. Note, that glioblastoma cell lines used in this work were biological samples from the previous study^23^.

##### TMT-labeling

Samples were resuspended in 100 mM HEPES (pH 8.5) and then labeled with the TMT10-plex kit (Thermo Fisher Scientific, San Jose, CA, USA) in accordance with the manufacturer’s instructions. The control samples were labeled with TMT-127N, TMT-127C, TMT-128N, and TMT-128C tags; and IFN treated samples with TMT-129N, TMT-129C, TMT-130N, and TMT-130C tags. Small quantities of all labeled samples were pooled, desalted, and analyzed by LC−MS/MS to evaluate the labeling efficiency. The quantities of peptides in each labeling channel were adjusted to the same average concentration before mixing.

##### Fractionation

Prior fractionation on a high pH system the TMT-labeled samples were desalted using cartridges for solid-phase extraction (Oasis HLB, 1 cc, 30 mg, 30 µm particle size, Waters, Milfold, CT, USA) and dried. Then samples were dissolved in solvent A (20 mM ammonium formate, pH 9.2). Peptides were separated on an ACQUITY UPLC M-class, CSHTM C18, 130, 1.5m, 300 mm x 100 mm column (Waters), using a Dionex Ultimate 3000 LC system (Thermo Fisher Scientific) equipped with a fraction collector. Fractions were concatenated into 10 and then dried.

#### *E. coli* with UPS1

*E. coli* cell pellet was resuspended in lysis buffer (100 mM tetraethylammonium bromide (TEAB), 5% Sodium dodecyl sulfate (SDS), PhosSTOP (Roche), and cOmplete protease inhibitor cocktail (Roche)).The mixture was then subjected to tip sonication on ice using a Bandelin Sonopuls HD2070 (Bandelin Electronic, Berlin, Germany) ultrasonic homogenizer (1 s on, 1 s off for 3 min with an amplitude of 40% and 3 min with amplitude of 70%). The sample rested for 30 min at room temperature following centrifugation for 15 min at 20000 g, supernatant was taken for following analysis. The digest was prepared by PAC protocol^24^. Briefly, the sample was diluted with ACN to a final concentration of 70% v/v and magnetic beads (MagReSyn HILIC) were added to the sample in mass ratio 2:1 beads:protein. Sample incubated for 30 min with occasional shaking. The sample was placed on the magnet and washed shortly with 100% ACN (twice) and with 70% EtOH for 10 sec. Next, the sample was removed from the magnet and dried gently in the fume hood. The sample was resuspended in 300 uL of 50 mM TEAB, reduction and alkylation was performed with 5 mM DTT for 30 min and 20 mM chloroacetamide (CAA) for 30 min, respectively. LysC (1:100) was added for 2h at 37 °C, and the sample was incubated with trypsin (1:100) overnight at 37 °C. Digestion was terminated with trifluoroacetic acid (TFA), after that sample was desalted and dried.

Universal Proteomics Standard-1 (UPS1) set (Sigma) containing 48 human proteins in equimolar concentrations of 5 pmol each was resuspended in 50 uL 50 mM TEAB buffer, then Disulfide bridges were reduced with 5 mM tris(2-carboxyethyl)phosphine (TCEP) and alkylated with 20 mM CAA for 30 min. LysC was added in ratio 1:100 for 2h at 37 °C, then trypsin was added in ratio 1:100 and the sample incubated overnight at 37 °C. UPS1 peptides were diluted serially in *E. coli* peptides to obtain 8 different UPS1 concentrations: 50, 25, 10, 5, 2.5, 1, 0.25, and 0.1 fmol of UPS1 per 1 μg of *E. coli* digest.

#### Data Acquisition

##### LC-MS1-only

The ultra-fast LC-MS1-only analysis of proteomic samples in this work was performed using Orbitrap Fusion Lumos mass spectrometer (Thermo Scientific, San Jose, CA, USA) coupled with UltiMate 3000 LC system (Thermo Fisher Scientific, Germering, Germany) and equipped with FAIMS Pro interface. Short LC gradient method was applied as reported previously^6^. Trap column µ-Precolumn C18 PepMap100 (5 µm, 300 µm, i.d. 5 mm, 100 Å) (Thermo Fisher Scientific, USA) and self-packed analytical column (Inertsil 3 µm, 75 µm i.d., 5 cm length) were employed for separation. Mobile phases were as follows: (A) 0.1 % formic acid (FA) in water; (B) 80 % ACN, 0.1 % FA in water. Loading solvent was 0.05 % TFA in water. The gradient was from 5 % to 35 % phase B in 4.8 min at 1.5 µL/min. Total method time including column washing and equilibration was 7.3 min. Field asymmetric ion mobility spectrometry (FAIMS) separations were performed with the following settings: inner and outer electrode temperatures were 100 °C; FAIMS carrier gas flow was 4.7 L/min; asymmetric waveform dispersion voltage (DV) was −5000 V; entrance plate voltage was 250 V. Compensation voltages (CV) -50 V, -65 V, and -80 V were used in a stepwise mode during LC-MS analysis. Mass spectrometry measurements were performed in MS1-only mode of acquisition. Full MS scans were acquired in a range from m/z 375 to 1500 at a resolution of 120 000 at *m/z* 200 with Automatic Gain Control (AGC) target of 4·10^5^, 1 microscan and 50 ms maximum injection time. Two hundred nanogram of glioblastoma digest and 1 ug of UPS-*E. coli* digest were loaded per injection.

##### DDA LFQ and TMT

MS/MS-based analysis of glioblastoma samples was performed using Q-Exactive HF mass spectrometer (Thermo Fisher Scientific, San Jose, CA, USA). Experimental label-free data for DDA mode were published elsewhere^23^. Experimental data for TMT were acquired in two ways: single-shot LC-MS/MS runs and deep proteome analysis with sample pre-fractionation. Trap column µ-Precolumn C18 PepMap100 (5 µm, 300 µm, i.d. 5 mm, 100 Å) (Thermo Fisher Scientific, USA) and self-packed analytical column (Inertsil 3 µm, 75 µm i.d., 25 cm length) were employed for separation. Mobile phases were as follows: (A) 0.1 % FA in water; (B) 95 % ACN, 0.1 % FA in water. The gradient was from 2 % to 5 % phase B in 1 min, to 33% B in 36 min, then to 45% B in 2 min at a flow rate 300 nL/min. Total method time including column washing and equilibration was 62 min. Data was acquired in top15 mode. Full MS scans were acquired in a range from *m/z* 350 to 1400 at a resolution of 120 000 at *m/z* 200 with AGC target of 3e6, 1 microscan and 50 ms maximum injection time. For higher energy collision-induced dissociation (HCD) MS/MS scans, the normalized collision energy was set to 32, and the resolution was 60 000 at *m/z* 200. Precursor ions were isolated in a 1.2 Th window and accumulated for a maximum of 110 ms or until the AGC target of 2e5 charges was reached. Precursors of charge states from 2+ to 6+ (inclusive) were scheduled for fragmentation. Previously targeted precursors were dynamically excluded from fragmentation for 15 s. One microgram of glioblastoma cell line samples were loaded on the column for single-shot and fractionated samples.

##### DIA

Experimental data for the UPS-*E. coli* mixture recorded in DIA mode are published elsewhere^25^.

#### Data Analysis

Raw files were converted into mzML format using ThermoRawFileParser^26^ (v. 1.3.4).

##### DirectMS1

Peptide features were detected using biosaur2 (v. 0.1.7) which is an update of previously developed Biosaur^27^ software for feature detection. biosaur2 is freely available at https://github.com/markmipt/biosaur2 under Apache 2.0 license. The difference between biosaur2 and original Biosaur is discussed in Supporting materials. Data was analyzed by using core DirectMS1’s search engine, ms1searchpy (v. 2.3.7), freely available at https://github.com/markmipt/ms1searchpy under Apache 2.0 license. Parameters for the search were as follows: minimum 1 scan for detected peptide isotopic cluster; minimum one visible ^13^C isotope; charge states from 1+ to 6+, no missed cleavage sites, carbamidomethylation of cysteine as a fixed modification and 8 ppm initial mass accuracy, and peptide length range of 6 to 30 amino acid residues.

DirectMS1’s search workflow was improved in this work (see details on updates in the Supporting materials) and now provides more than 2500 proteins at 1% FDR for 5 min LC-FAIMS/DirectMS1 data obtained earlier for 200 ng of HeLa using Orbitrap Lumos instrument^7^. The boost also came from employing an updated version 1.1.2 of peptide retention time algorithm DeepLC^28^.

##### MS/MS-based data

MS/MS analyses for both DDA LFQ and TMT-labed data were performed using ProteomeDiscoverer (v. 2.5) with Sequest HT search engine. Parameters for the search were as follows: up to two missed cleavage sites, carbamidomethylation of cysteine as fixed modification, oxidation of methionine as variable modification, 10 ppm, and 0.05 Da precursor and fragment mass accuracies. Additionally, DDA data was analyzed by using a combination of IdentiPy^29^ (v. 0.3.5) search engine and postsearch tool Scavager^30^ (v. 0.2.9). Diffacto^31^ (v. 1.0.6) was used for protein quantitation with a global normalization strategy on median values.

DIA data was analyzed with Spectronaut^32^ (v. 15.2.210819.50606) using database searching with following parameters: peptide length from 7 to 30, maximum missed cleavages 2 with trypsin cleavage rule, carbamidomethylation of cysteines was set as fix modification and oxidation of methionine as variable, mass tolerance strategy was dynamic, MS1 and MS2 correction factor was set to 1. Search results filtered to 1% FDR and quantitation results filtered to 5% FDR. QUANT 2.0 was used for protein quantitation with quantity MS2-level and area quantity type, and global normalization strategy on median values.

Searches were performed against combinations of Swiss-Prot human (20241 protein sequence), *E. coli* (4431 proteins) or UPS (48 proteins). Decoy proteins were created by built-in methods within each software. A special decoy generation was implemented in ms1searchpy and discussed in detail in Supporting materials.

#### Data Availability

The data sets generated and analyzed during the current study have been deposited to the ProteomeXchange Consortium via the PRIDE partner repositories with the data set identifiers PXD033350 and PXD033355. Additionally, the data from repositories PXD026600 and PXD022906 were used in this study. Supplementary Table S1 provides a list and the annotations for all experimental data used in this work.

## Results and Discussion

### DirectMS1Quant: workflow of the method

Most popular proteomics quantitation methods take into account the number of identified peptides^33,34^, intensities of peaks of peptide ions in MS1 spectra^20,22,31,35^, fragment ion (Sin) or TMT label intensities in MS2 or MS3 spectra. Note that these methods typically employ the information from DDA LFQ, DDA TMT, or DIA identifications which have low peptide FDR, and, thus, they are not utilizing for the presence of false positive identifications while performing quantitation analysis. On the contrary, DirectMS1 results contain protein and peptide identifications at specified low and high uncontrolled FDR levels, respectively. The latter depends on the search parameters, database and the experimental data itself. For example, we estimated the peptide-level FDR of 20 to 30% for 5 min LC-FAIMS/DirectMS1 data obtained earlier for 200 ng of HeLa using Orbitrap Lumos instrument^7^. This peptide FDR level was estimated for 2000 top scored proteins at 1% protein-level FDR. Moreover, the FDR level at peptide feature matches (PFMs) for all proteins before filtering was as high as 90% on average. These enormously high numbers of false matches require a special treatment during quantitation analysis implemented here in DirectMS1Quant. We propose the quantitation workflow, which is schematically shown in Figure 1. The input of the software presents the ms1searchpy results grouped by samples. At first step, it combines all identified target proteins into a single list of identified proteins for all files. Next, it extends that list with proteins which share at least one matched peptide with the proteins in the list. In this way, we add all isoforms and homologous proteins to the quantitation procedure. At this step, the protein list is also extended with decoy versions of target proteins. At the third step, DirectMS1Quant extracts all PFMs which belong to the above extended list of proteins from MS1 spectra. There are multiple cases when a peptide sequence is matched to multiple PFMs which leads to a problem of multiple potential intensity values for this peptide. To solve this problem, all PFMs are sorted by intensity and only the maximum intensity is assigned to the particular peptide’s multidimensional vector, which is defined as the combination of sequence, charge state, and FAIMS CV value. In this way, a peptide sequence, e.g. identified at two different charge states will be used twice for the quantitation purpose, etc.. The peptide vectors which are not identified in some of the experimental files have missing intensities values in these files. At the next step, all peptides with the number of missing values exceeding the user-defined threshold (*min_samples* parameter) are excluded from the subsequent analysis. By default, DirectMS1Quant sets that value to half of the number of the experimental files. Then, all peptide intensities are normalized by datafile-specific median intensities, and all missing values are replaced by datafile-specific minimal intensities. The above-mentioned median and minimal intensities are the ones of extracted peptide vectors. The list of all identified peptide sequences is further analyzed, including the calculation of the *p*-value of the Independent Samples *t*-test and log2 fold change between the average intensities of two samples for each peptide. All peptides with *p*-value less than 0.05 and absolute value of fold change higher than a user-defined threshold are marked as significantly changed (“significant peptides”). By default, the fold change threshold is 3 standard deviations from the distribution of background calculated using peptides with *p*-values higher than 0.05. Note that these *p*-values are used as the raw numbers with no correction applied. At the next step, the total number of peptides passed all quantitation filters, *n*, and the number of co-directed significant peptides by fold change, *k*, are calculated for each protein. The latter is the maximal value of significant peptides with the same sign of log2 fold change values. For example, if a protein has 10 peptide matches passed filters, in which 2 and 5 of them are significant with negative and positive fold changes, respectively, the above two numbers will be *n*=10 and *k*=5.

**Figure 1.**
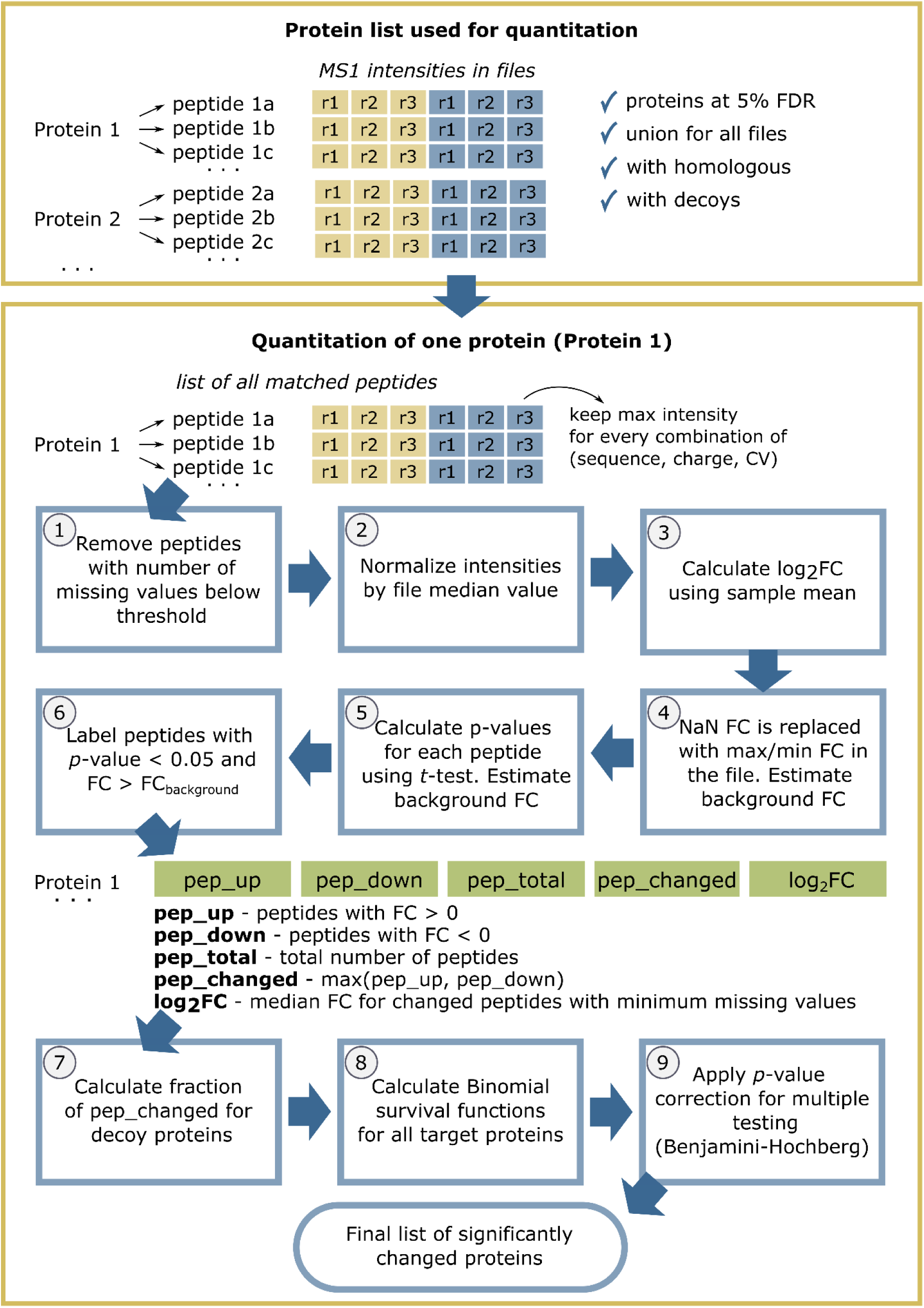
DirectMS1Quant workflow.

Now, recall that we have decoy proteins in the search database. Taking these proteins separately, the algorithm calculates the probability of a peptide to be significant by chance as follows:

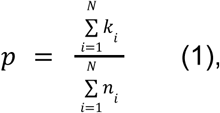

in which N is the number of decoy proteins.

After calculation of this probability, all decoy peptides and proteins are not used anymore for estimating the quantitation FDR. Also, the proteins having no significant peptides are excluded from the analysis.

Finally, the algorithm performs quantitation statistics analysis using binomial distribution function. Here, a protein is described by binomial distribution with the number of successful matches, *k*, (the number of co-directed significant peptides) out of the number of all identified peptides passed all quantitation filters, *n*, (each considered as an independent experiment) with success probability *p*, calculated using Eq.1. DirectMS1Quant uses the cumulative distribution function of the binomial distribution to calculate the probability of a protein to have *k* or more randomly co-directed significant peptides. Probability is set to 1 for all proteins which have only 1 significant peptide. This is similar to the “one-hit wonder” exclusion in protein identification. Calculated probabilities are corrected using Benjamini-Hochberg procedure. All proteins with probability less than 0.05 are reported as significantly changed with fold change estimated as median value of fold changes of co-directed significant peptides with minimum number of missing values. For example, if a protein has 10 peptide matches, 5 of them are significantly changed and only 3 of these 5 peptides have no missing values, then protein fold change will be estimated using these 3 peptides only.

DirectMS1Quant reports two fold changes for proteins calculated using either normalized or raw peptide intensities.

### UPS-*E. coli* data: quantitation results for the characterized sample

Five min LC-MS1-only data for DirectMS1 analysis was acquired as described in the methods section. As a benchmark of the quality of quantitation, we used the 90-min DIA dataset obtained recently^25^. In the latter report, the authors evaluated an extensive list of DIA data processing software and performed sample analysis using four different DIA acquisition schemes and two types of searches, library-based and library-free. Reportedly, only Spectronaut^32^ software successfully provided stable results for all combinations of acquisition schemes and types of searches with the most efficient combination of library-free search and the narrow window DIA acquisition. In this work we re-analyzed those data to obtain the Spectronaut’s quantitation results using the original search parameters and the built-in Spectronaut quantitation module.

The searches for both Spectronaut and DirectMS1 methods were performed against a combined *E. coli*-UPS protein database. Separately, we performed DirectMS1 analysis using full Swiss-Prot Human-*E. coli* database (see Supporting materials).

DirectMS1 reported from 1 to 46 UPS proteins at 1% FDR for 0.1 to 50 fmol concentration ranges, respectively (see Supporting materials), as well as ca. 1200 *E. coli* proteins on average per experimental run. On the other hand, Spectronaut reported 48 UPS proteins and ca. 1900 *E. coli* proteins in the aggregated results for all data without separation of the samples by UPS concentration ranges that makes direct comparison quite dubious. However, the efficiency of quantitation is different from the identification efficiency. This is further demonstrated in Figure 2a by the number of UPS proteins whose changes are reported as statistically significant for Spectronaut and DirectMS1Quant methods. It shows that DIA reported more UPS proteins for 25 and 50 fmol samples compared with lower concentration ones. In the DirectMS1Quant method a number of proteins are missing among the identifications even for the samples with high concentrations of spiked UPS proteins. At low concentrations of spiked UPS proteins, 2.5 and 5 fmol samples, DirectMS1Quant starts reporting a larger number of correctly quantified UPS proteins compared with DIA-Spectronaut results. These results are more important than the results for UPS proteins spiked at higher concentrations because 2.5 and 5 fmol samples represent protein concentrations close to the ones in the typical biological samples. Indeed, there are approximately 5000 proteins being typically identified in the standard hour-long MS/MS-based DDA proteome analysis of 200 ng of human cell line. This may be directly translated into the average concentration of proteins in such sample close 1-2 fmol. Contrary to the UPS proteins, reporting *E. coli* proteins as significantly changed demonstrates the deficiencies of one or the other quantitation methods. Figure 2b shows the number of “false” *E. coli* proteins reported as significantly changed. To benchmark the sensitivity and specificity of DirectMS1Quant against Spectronaut, we calculated the total number of *E. coli* and UPS proteins reported as significant across all 28 comparisons. The upper limit for the number of reported UPS proteins is 1344, which is 28 comparisons multiplied by 48 proteins in the UPS standard. For all 28 comparisons, DirectMS1Quant and Spectronaut reported 35 and 94 such *E. coli* proteins compared with 868 and 807 UPS proteins, respectively. Thus, DirectMS1Quant and Spectronaut had the estimated empirical quantitation FDRs of 4 and 12% on average, while the FDR requirement settings were 5% for both methods.

**Figure 2.**
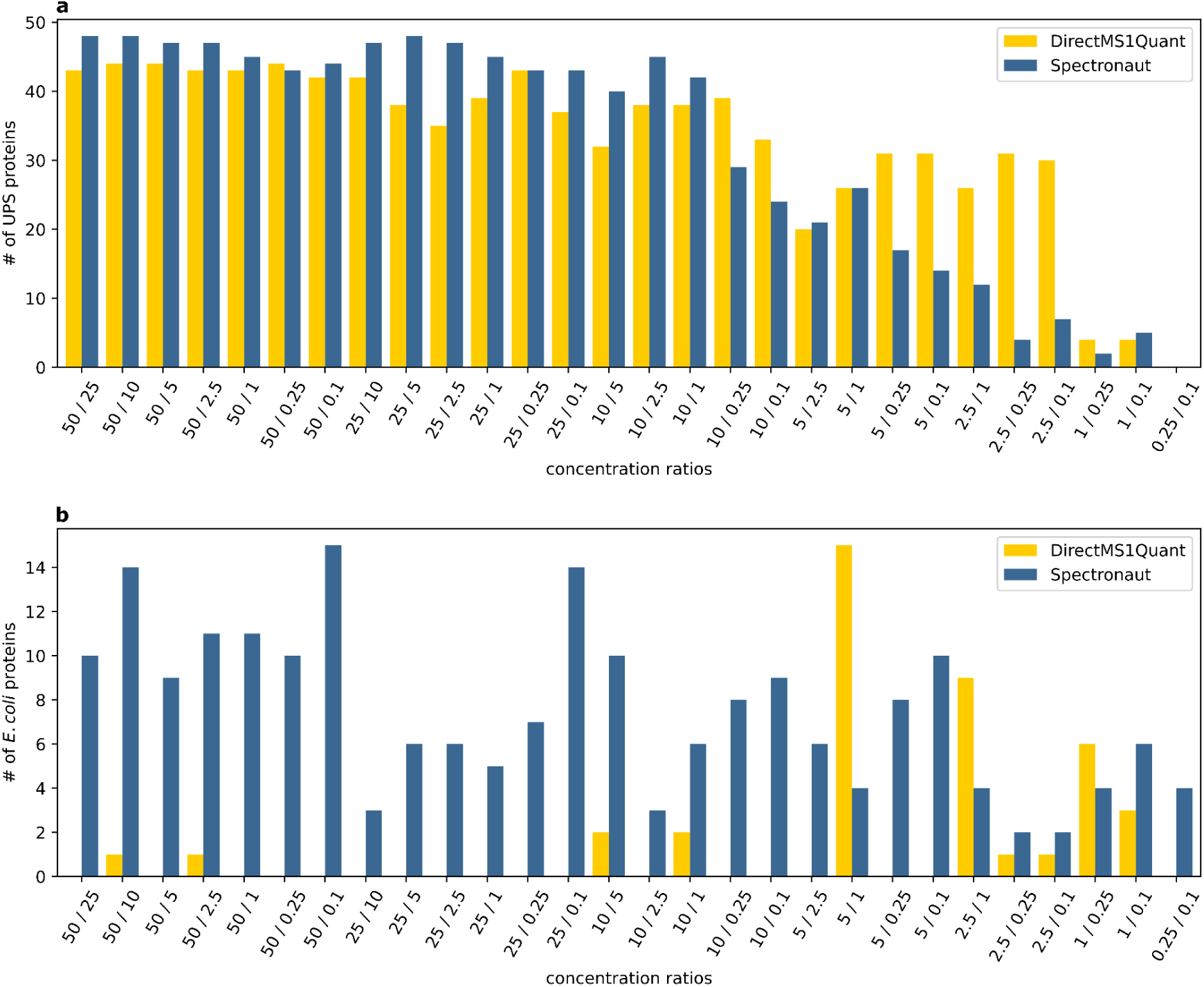
The number of significantly changed (a) UPS and (b) *E. coli* proteins reported at 5% quantitation FDR by DirectMS1Quant and Spectronaut analyses using *E. coli*+UPS protein databases for all 28 pairwise concentration comparisons.

Notably, the DirectMS1Quant method delivers similar quantitation results regardless of the FDR threshold selected for protein identifications (see Supporting materials). For consistency, we employed 5% identification and 5% quantitation FDR thresholds here for all data analyzed in this work.

An effect of missing values on DirectMS1Quant method is discussed in Supporting materials.

Additionally, DirectMS1Quant method was compared with Diffacto which was the only available method for quantitation analysis of DirectMS1 results (see Supporting materials).

Then, we estimated the quantitation accuracy of the method as the root mean squared errors calculated using the following equation:

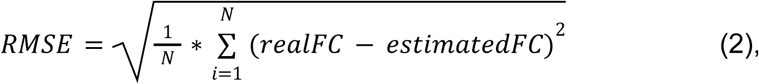

in which N is the number of UPS proteins reported by the method for a given UPS concentration sample, realFC and estimatedFC are experimental and reported by software protein fold changes in log_2_ scale.

The calculated RMSE values for DirectMS1Quant and Spectronaut are shown in Figure 3. Note that the DirectMS1Quant values were calculated using both normalized (“normalized”) and raw (“raw”) peptide intensities. The average RMSE values of 1.5, 1.51 and 1.8 were obtained for all 28 comparisons of different UPS concentration samples for DirectMS1Quant “normalized”, DirectMS1Quant “raw” and Spectronaut methods, respectively.

**Figure 3.**
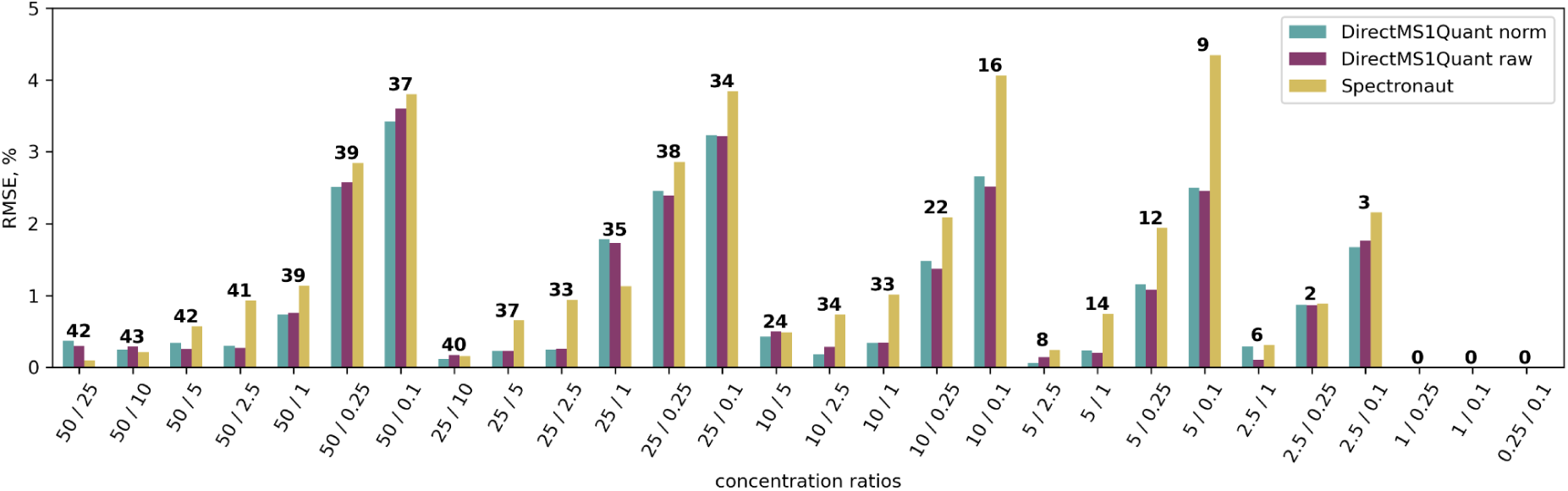
RMSE values for UPS proteins reported as significantly changed by both DirectMS1Quant and Spectronaut methods. Values above bins show the number of UPS proteins used for RMSE calculation.

### Glioblastoma data

To explore the DirectMS1Quant sensitivity for quantitative description of cellular response to stimuli, two glioblastoma, GBM3821 and GBM4114, and the astrocyte (AN) cells treated with interferon alpha (IFN*α*) were used. Earlier it has been demonstrated that AN and GBM3821 cells respond to IFN*α* by upregulation of dozens of IFN-induced proteins, while GBM4114 has impaired interferon-dependent antiviral mechanisms and does not respond to IFN stimulation^23^. In this work, IFN response of these cellular models was quantified using different workflows shown in **Table 1**. The workflows included quantitation of label free data obtained using 2h LC gradients, TMT-labeled data collected using high pH fractionation (10 fractions, 40 min LC gradient each), single-shot 40 min LC gradient in DDA mode, as well as data acquired by 5 min DirectMS1 method. We focused our analysis on IFN-stimulated genes that are expected to be regulated only in AN and GBM3821 cell lines. As shown in the table for the results of quantitative proteome profiling, DirectMS1Quant confirms previous results and reports 13 upregulated genes in both GBM3821 and AN cells, which were assigned to IFN-stimulated genes. Our negative control, GBM4114 cells, did not respond by any changes in IFN-stimulated proteins with either of the experimental workflows compared in this study. These results are comparable with the ones obtained using community standard TMT analyses (high pH prefractionation) and significantly outperform 40 min single-shot TMT analysis, which provides the same total instrumentation time as the 5 min LC-DirectMS1. Long gradient DDA LFQ data provided the highest number of regulated IFN-stimulated gene products but it takes 40-fold more instrumentation time compared to DirectMS1 or single-shot TMT analyses.

**Table 1.** Summary of the proteins significantly upregulated in glioblastoma multiforme (GBM) and astrocyte (AN) cells in response to 100 U/mL IFN*α* treatment across different quantitative proteome analysis workflows. Differentially expressed proteins were selected according to the following criteria: |log_2_FC| >= 1 standard deviations of background fold changes, fdr_bh <= 0.05. IFN-stimulated gene products were recognized using public databases^23,36,37^. Designations: PD - Proteome Discoverer v.2.5; NSAF - normalized spectral abundance factor; Id,Scav - Identipy and Scavager; Diff - Diffacto; DiMS1Q - DirectMS1Quant.

| Cells | IFN response established elsewhere<sup>23</sup> | Number of IFN-stimulated / total genes coding upregulated proteins observed in this study, |  |  |  |  |  |  |  |  |
| --- | --- | --- | --- | --- | --- | --- | --- | --- | --- | --- |
|  |  | 2 h LC gradient, LFQ, DDA, no fractionation (total 5 days) |  |  | 40 min LC gradient, TMT10plex, DDA, 10 fractions (total 30 h) |  | 40 min LC gradient, TMT10plex, DDA, no fractionation (total 3 h) |  | 5 min LC-MS1 (total 3 h) |  |
|  |  | PD, Diff | Id,Scav,Diff | Id,Scav,NSAF | PD, Diff | Id,Scav,Diff | PD, Diff | Id,Scav,Diff | Diff | DiMS1Q |
| GBM4114 | Impaired JAK/STAT signal: No upregulation of IFN-induced proteins | 0 / 0 | 1 / 22 | 0 / 0 | 0 / 0 | 0 / 0 | 0 / 0 | 0 / 0 | 0 / 0 | 0 / 0 |
| GBM3821 | Active JAK/STAT signal: 50-70 IFN upregulated proteins | 52 / 76 | 8 / 22 | 22 / 34 | 14 / 17 | 8 / 10 | 6 / 7 | 2 / 2 | 9 / 10 | 13 / 13 |
| AN |  | 52 / 76 | 11 / 32 | 26 / 54 | 35 / 38 | 10 / 17 | 6 / 7 | 2 / 2 | 16 / 18 | 13 / 13 |

To further demonstrate the quantitative capabilities of ultrafast DirectMS1 method, ranking of interferon pathway functionality was performed at the level of gene ontology enrichments (**Figure 4**). Processes were ranked by GO_score^38^ calculated using the following equation:

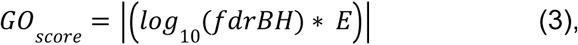

where fdr_BH is the Benjamini-Hochberg false discovery rate and E is the enrichment, both calculated by STRING db.

**Figure 4.**
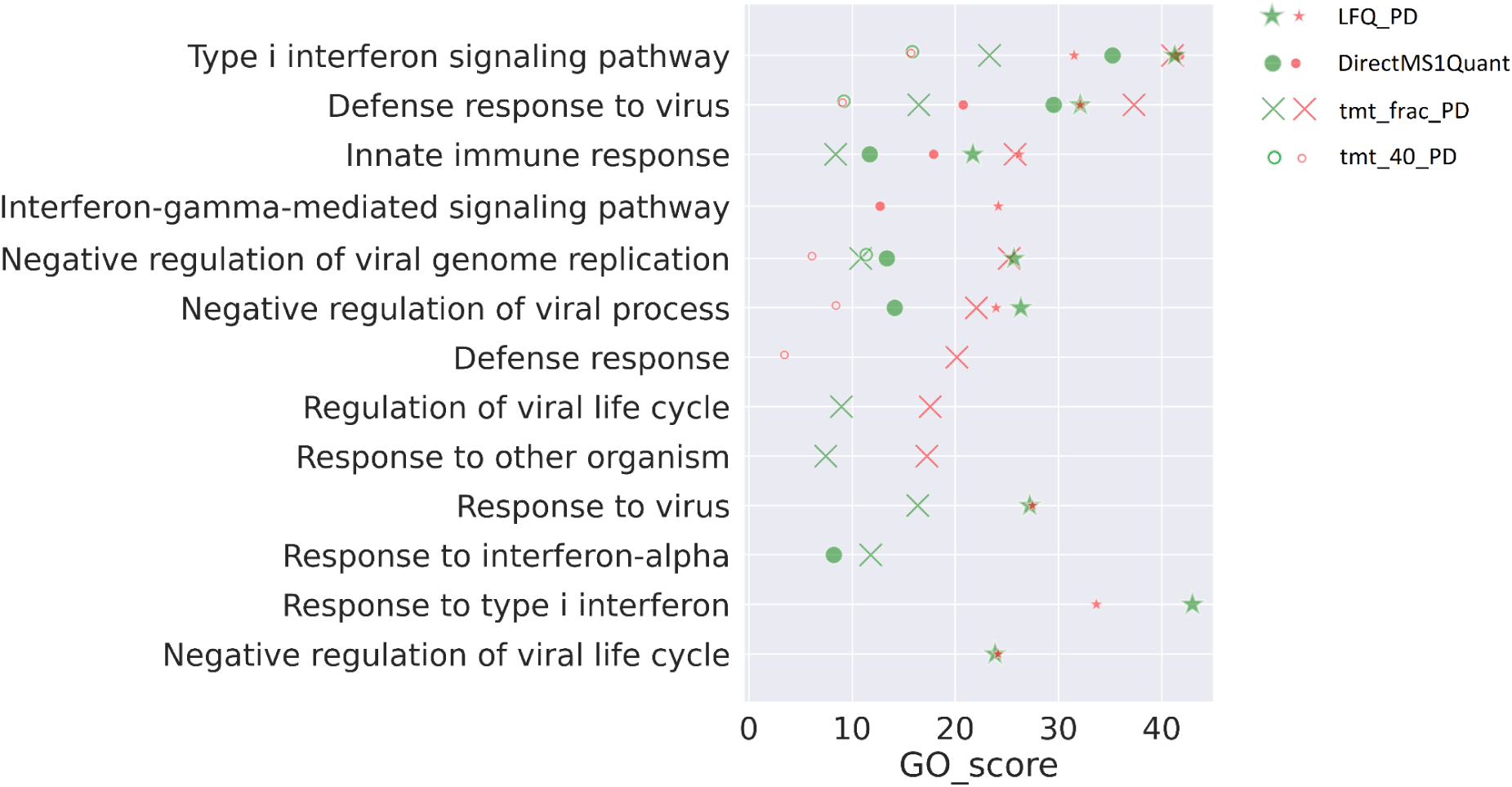
Top-8 enriched biological processes in GBM3821 (green markers) and AN (red markers) cells in response to IFN treatment revealed by different quantitation workflows. Quantification was performed using Proteome Discoverer (PD) and DirectMS1Quant. Only processes identified by at least two quantitation methods are shown.

Analysis of biological processes in IFN-stimulated GBM3821 (green markers) and AN (red markers) cells has revealed the type I interferon signaling pathway as most confident and enriched. Note, that this pathway was correctly identified by all quantitation workflows compared. Since GBM4114 cells did not respond to IFN stimulation, no enriched interferon-related pathways were reconstructed with any quantitation workflows and their GO_score can be attributed to zero. Sensitivity and selectivity of DirectMS1Quant post-processing of 5 min LC-MS1 data was enough to score the pathway enrichment and functionality close to the results of label-free and TMT-based quantitation on fractionated samples obtained with ProteomeDiscoverer. However, ultrafast LC separation with MS1 data acquisition provided at least 10-fold gain in instrument time compared with the analysis of fractionated TMT-labeled samples, not even mentioning 2 hour DDA LFQ results which were obtained after five days of experiments.

## Conclusions

The results obtained in this study have shown that the ultrafast DirectMS1 method provides proteome-wide quantitation performance comparable with the current community standard MS/MS-based approaches including DIA and TMT-based multiplexed DDA proteomics. However, DirectMS1 allows reducing the instrumentation time to perform the proteome-wide analyses 10-fold compared with, e.g. 1 hour long TMT 10plex DDA with sample pre-fractionation. The method also alleviates the experimental complexity needed for the quantitative proteome characterization as being label- and MS/MS-free. The DirectMS1Quant quantitation algorithm developed in this work and integrated in the DirectMS1 method has been tested using samples of different types and complexity including the assessment of pathway functionality in the expression proteomics applications.

## Software availability

DirectMS1Quant software is freely available under Apache 2.0 license and can be installed along with ms1searchpy from https://github.com/markmipt/ms1searchpy under Apache 2.0 license.

## ACKNOWLEDGMENTS

UPS-*E. coli* and Glioblastoma samples proteomic analyses, as well as methods and software development were performed with financial support from the Russian Science Foundation, grant no. 21-74-10128 to M.V.I. Proteomics and mass spectrometry research at the University of Southern Denmark SDU is supported by generous grants to the VILLUM Center for Bioanalytical Sciences (VILLUM Foundation grant no. 7292) and PRO-MS: Danish National Mass Spectrometry Platform for Functional Proteomics (grant no. 5072-00007B).

