## Supporting materials for "DirectMS1Quant: ultrafast quantitative proteomics with MS/MS-free mass spectrometry"

### Decoy database generation

Generation of decoy database is one of the key steps in DirectMS1Quant search pipeline as it critically affects the method efficiency. Note that the previous version of the DirectMS1 method’s software implementation employed a protein sequence shuffling approach for decoys. However, this widely-used approach has a drawback for the method as it results in a higher number of unique decoy peptides compared to the number of unique target peptides due to the presence of homologous target proteins[1]. This does not create a problem for protein FDR estimation because the number of theoretical peptides are taken into account in the calculation of protein scores. On the other hand, the lower number of shared decoy peptides results in more conservative FDR estimation due to the “one feature - one peptide - one protein” rule used also in the protein scoring. Indeed, some of the peptides of a target protein will go to the other high scored proteins. This will create a bias towards decoy proteins, which will receive higher scores due to more random PFMs matching their sequences. This problem was solved in the DirectMS1 method by using a modified pseudo shuffle decoy generation method. Pseudo shuffle method is identical to the shuffle method but shuffles only parts of the target protein sequences between cleavage sites (R and K in case of trypsin). To solve the difference between the number of unique target and decoy peptides, which may occur due to protein homology, we use one and same decoy peptides for target peptides of the same sequence from different homologous proteins.

### Feature detection in MS1 spectra: Biosaur vs biosaur2

In this study we used a rewritten version of Biosaur - our previously published software for peptide feature detection. While we made minor changes in the software logic and algorithm, rewritten code contains less bugs and is more stable. For example, biosaur2 can report peptide isotopic clusters which were detected only in a single MS1 scan. The latter increases the sensitivity of DirectMS1 analysis. Current version of DirectMS1 supports three feature detection algorithms which are Dinosaur, Biosaur and biosaur2. Biosaur and biosaur2 were compared using previously published data[2] of 200 ng of HeLa analyzed used Orbitrap Lumos with 3 CV FAIMS values. Dinosaur does not support FAIMS data and was not used in that comparison. The average number of detected peptide features are 100500 and 179500 for

Biosaur and biosaur2, respectively. The average number of identified proteins using DirectMS1 at 1% FDR are 2202 and 2509 for Biosaur and biosaur2, respectively.

**Mass recalibration**

DirectMS1Quant performs mass re-calibration before the final search, which accounts for FAIMS data. Upon re-calibration, the mass differences between experimental and theoretical masses of peptides, assigned to high score protein sequences after preliminary search (see the DirectMS1 search workflow description in the earlier work[3], are grouped by FAIMS CVs. Each of the groups is further divided into 5 subgroups according to the percentile of retention time range. Then, the median mass error for a group is calculated and subtracted from the mass error for each peptide within the group. Mass re-calibration as described provides an increase albeit insignificant in the number of identified proteins for the data used in this work.

**Machine learning based workflow implemented in DirectMS1 method**

A search engine of DirectMS1method, ms1searchpy, employs LightGBM machine learning package[4] for estimation of peptide-feature-matches (PFMs) quality. The machine learning workflow described earlier[2] performed division of all matches into target and decoy peptide groups and the model was trained to predict which group a particular PFM belongs to. The new version of ms1searchpy splits PFMs into two groups of target PFMs from proteins below and above 25% FDR threshold instead of target and decoy group splitting. The described modification of the search algorithm better assists in estimating the quality of PFM matching by reducing the risk of prediction model overfitting as it relies on the differences between true and false target matches instead of the differences between target and decoy matches.

**The effect of the search space**

To evaluate the real quantitative FDR provided by the method the 5-min DirectMS1 UPS-*E. coli* data were analyzed using two databases: combined whole Swiss-Prot Human/*E. coli* and truncated UPS/*E. coli*. When searched against the first one, the quantitation analysis by DirectMS1Quant has resulted in 805 UPS proteins in total reported as differentially expressed for all 28 pairwise concentration comparisons (Figure S1a). This number is slightly less (about 7% smaller) compared with 867 UPS proteins resulted from the searches against UPS-*E. coli* database. Note that quantitation of *E. coli* proteins constitutes the presence of false positive identifications and the average percentage of *E. coli* proteins among all identified proteins for UPS-*E. coli* protein database was 3.6%, which is favorably comparable with the expected quantitative FDR of 5% by default settings (Figure S1b). However, the searches against the whole Swiss-Prot Human-*E. coli* protein database have resulted in as much as 18.9% of quantified *E. coli* and human non-UPS proteins on average for all pairwise comparisons (Figure S1c). Upon a closer look at these identifications, we found that most of the reported non-UPS human proteins have at least one matched peptide shared with any of the UPS proteins or common contaminants (shown in green and blue in Figure S1c, respectively). We also found two human proteins, P05452 and P40926, without any relation to UPS, yet, continuously reported as differentially expressed for all UPS concentrations with similar fold changes (shown in blue in Figure S1c). These proteins were not found in either identification, nor quantitation results obtained for DIA data using DirectMSQuant and searches against the whole Swiss-Prot Human-*E. coli* database. One of the straightforward suggestions is that these proteins are really present in the UPS1 sample used in our work as contaminants. To validate this assumption, we performed a standard DDA MS/MS-based analysis for the UPS1 sample used and identified 9 and 11 peptides at 1% FDR for these suspicious proteins, respectively. After removing homologous and contaminant proteins from the quantitation results, the estimated empirical FDR for the 5-min DirectMS1 results obtained for UPS-*E. coli* sample using DirectMS1Quant method and search against the whole Swiss-Prot Human-*E. coli* database become 3.8% compared with expected FDR of 5%.

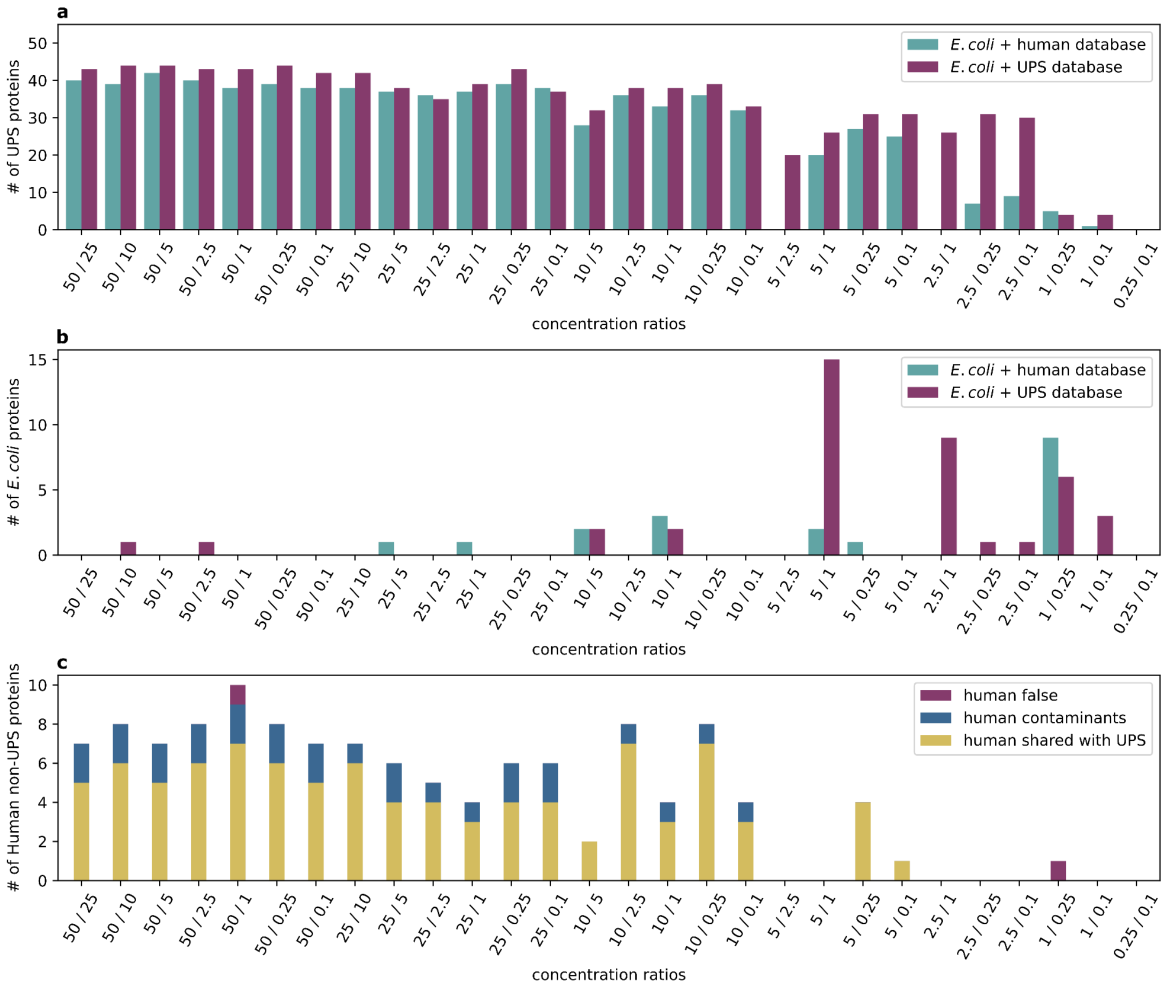


**Figure S1.** The number of (a) UPS, (b) *E. coli* and (c) human non-UPS proteins for DirectMS1Quant analysis using *E. coli*+UPS and *E. coli*+human protein databases.

### Effect of the identification FDR level setting on the quantitation

Identification analysis of UPS-*E. coli* data was done using two identification FDR thresholds: 1% and 5%. The average number of identified *E. coli* proteins were 1126 and 1395 for 1% and 5% FDR thresholds, respectively. The number of UPS proteins identified in all 3 replicates at 1% and 5% FDR are shown in Figure S2a. For comparison, we performed quantitation analysis for the identifications obtained using three different levels of FDR, including 1%, 5% and 100%. The latter is actually the same as using all proteins from the search database. To benchmark the sensitivity and specificity of DirectMS1Quant against Spectronaut, we calculated the total number of *E. coli* and UPS proteins reported as significant across all 28 comparisons. The upper limit for the number of reported UPS proteins is 1344, which is 28 comparisons multiplied by 48 proteins in the UPS standard. The numbers of reported differentially expressed UPS proteins were 874, 867 and 849 for 1, 5 and 100% identification FDR, respectively, for all 28 pairwise concentration comparisons. On the other hand, the numbers of reported *E. coli* proteins were 30, 32 and 22 for 1, 5 and 100% FDR levels, respectively. The details of these results for different spike-in UPS concentration and pairwise concentration comparisons are shown in Figure S2b. These results demonstrate that the DirectMS1Quant quantitation workflow filters effectively the false positives (Figure S2c).

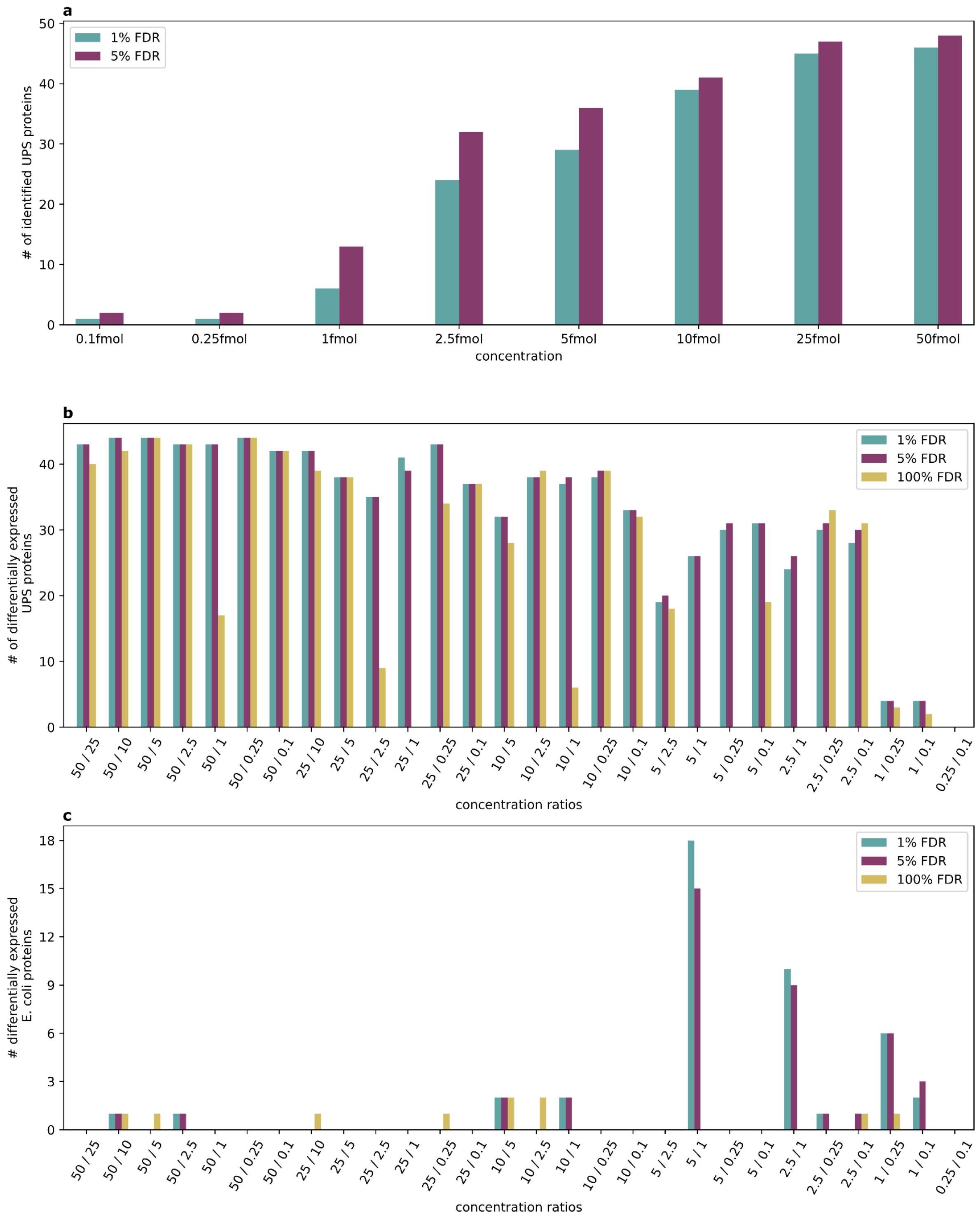


**Figure S2**. The number of (a) UPS proteins identified in all three replicates for 1% and 5% identification FDR, (b) differentially expressed UPS and (c) differentially expressed *E. coli*

proteins at 5% quantitation FDR for DirectMS1quant analysis using 1%, 5% and 100% identification FDR threshold. *E. coli*-UPS protein database was used for the identification.

**Dealing with missing proteins**

There is a special case of missing values when some of the proteins detected in one sample are being missed in the other one. For the UPS-*E. coli* data we performed multiple pairwise comparisons of two samples in three technical replicates each. Peptides completely missed in one sample will pass statistical tests when their missing values are imputed using the *min_samples* threshold set to 3 (half of the total 6 data files). On the other hand, these peptides will be removed from the analysis if the threshold is set to 4. Thus, we can roughly estimate the efficiency of our quantitation  in these particular cases by comparing the quantitation results for these two thresholds. DirectMS1Quant was used for UPS-*E. coli* data using 3 and 4 as the thresholds for the minimal number of samples, in which a particular peptide was identified (Figure S3). The results show no significant difference in both the quantitation accuracy and the sensitivity of the analysis for the UPS concentration ranges of 2.5 to 50 fmol. However, there is significant reduction in the number of reported differentially expressed UPS proteins for 1 fmol data and the threshold setting of 4 minimal samples. We explain this observation by missing some of the peptide-feature matches for part of the UPS proteins. The fraction of these missing values is expectedly even higher for 0.25 and 0.1 fmol UPS concentrations as further shown in Figure S3a. These results show that the imputation algorithm implemented in DirectMS1Quant correctly reports differentially expressed proteins completely missing in some samples, yet, not for the price of increased level of false identifications as shown in Figure S3b.

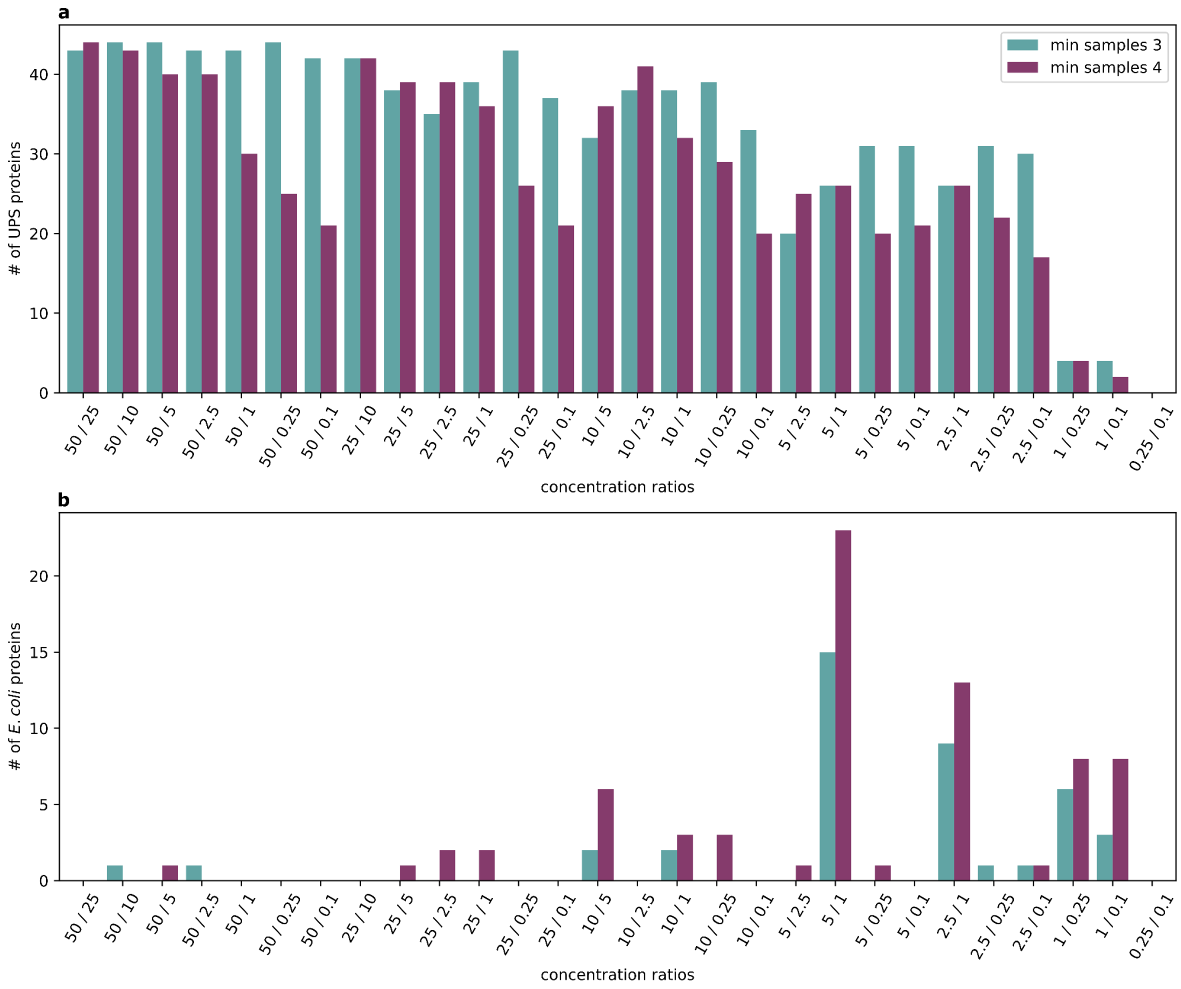

**Figure S3.** The number of differentially expressed UPS and *E. coli* proteins at 5% quantitation FDR for different minimal number of samples thresholds. Results are shown for DirectMS1Quant analyses using UPS-*E. coli* protein database.

**DirectMS1Quant vs Diffacto**. Quantitation analysis for DirectMS1 results was done using two different algorithms: DirectMS1Quant proposed in this manuscript and Diffacto which was the method of choice for quantitation analysis in our previous DirectMS1 manuscripts (see Figure S4). Diffacto provides a high number of false positives results when Benjamini-Hochberg correction was applied. Applying Bonferroni correction solves that issue but the sensitivity of analysis is lower compared to DirectMS1Quant results.

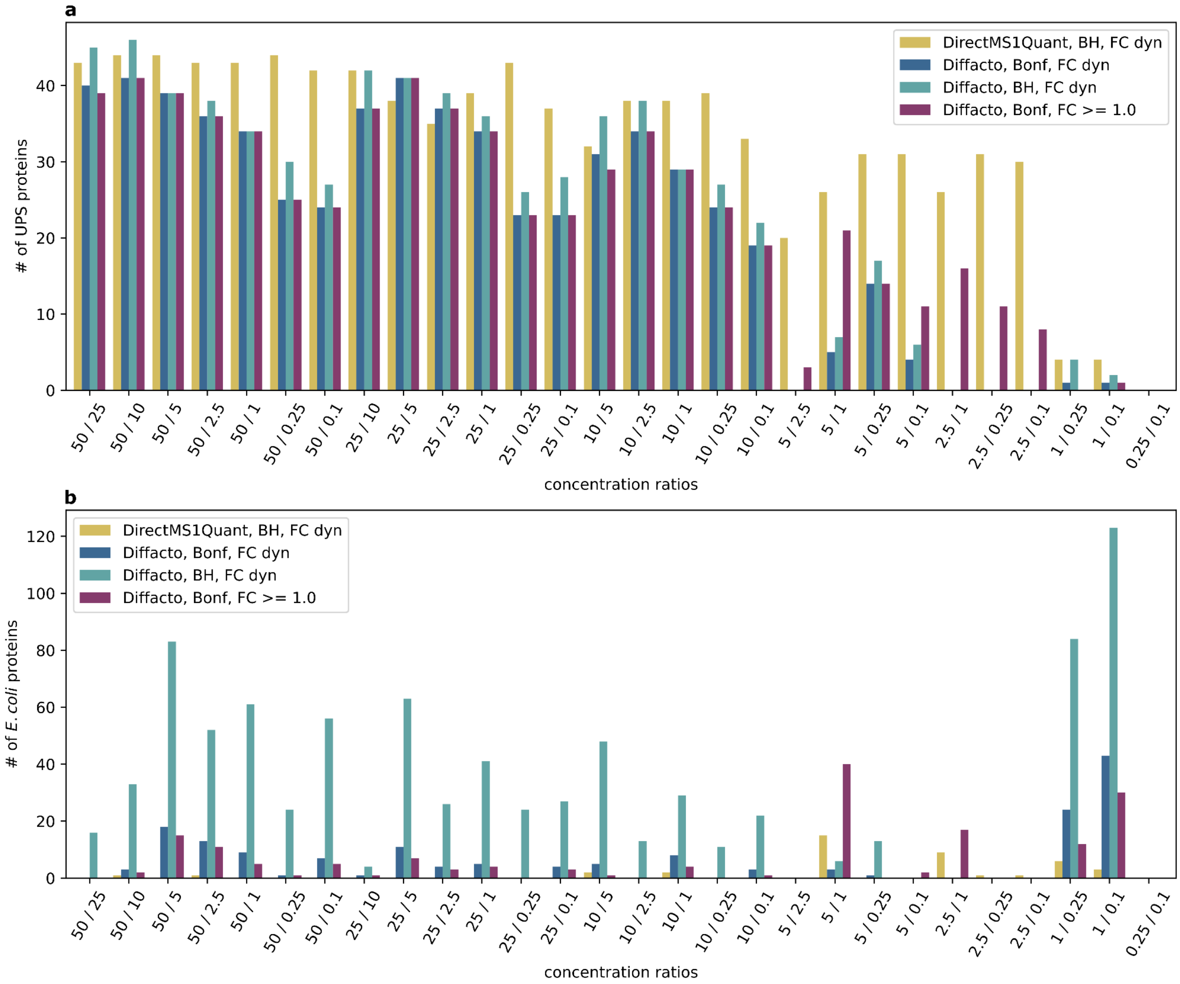


**Figure S4.** The number of differentially expressed UPS and *E. coli* proteins at 5% quantitation FDR for DirectMS1Quant and Diffacto analyses using UPS-*E. coli* protein database. Fold change thresholds were set either as dynamic estimation using background or fixed value. Correction of p-values was done using either Benjamini-Hochberg or more conservative Bonferroni correction.
